# Onset and stepwise extensions of recombination suppression are common in mating-type chromosomes of *Microbotryum* anther-smut fungi

**DOI:** 10.1101/2021.10.29.466223

**Authors:** Marine Duhamel, Fantin Carpentier, Dominik Begerow, Michael E. Hood, Ricardo C. Rodríguez de la Vega, Tatiana Giraud

## Abstract

Sex chromosomes and mating-type chromosomes can display large genomic regions without recombination. Recombination suppression often extended stepwise with time away from the sex- or mating-type-determining genes, generating evolutionary strata of differentiation between alternative sex or mating-type chromosomes. In anther-smut fungi of the *Microbotryum* genus, recombination suppression evolved repeatedly, linking the two mating-type loci and extended multiple times in regions distal to the mating-type genes. Here, we obtained high-quality genome assemblies of alternative mating types for four *Microbotryum* fungi. We found an additional event of independent chromosomal rearrangements bringing the two mating-type loci on the same chromosome followed by recombination suppression linking them. We also found, in a new clade analysed here, that recombination suppression between the two mating-type loci occurred in several steps, with first an ancestral recombination suppression between one of the mating-type locus and its centromere; later, completion of recombination suppression up to the second mating-type locus occurred independently in three species. The estimated dates of recombination suppression between the mating-type loci ranged from 0.15 to 3.58 million years ago. In total, this makes at least nine independent events of linkage between the mating-type loci across the *Microbotryum* genus. Several mating-type locus linkage events occurred through the same types of chromosomal rearrangements, where similar chromosome fissions at centromeres represent convergence in the genomic changes leading to the phenotypic convergence. These findings further highlight *Microbotryum* fungi as excellent models to study the evolution of recombination suppression.

## Introduction

Sex chromosomes in plants and animals can exhibit large genomic regions without recombination (Bachtrog et al., 2014; Bergero and Charlesworth, 2009). Recombination suppression often extended away from sex-determining genes in a stepwise manner, generating discrete strata of genetic differentiation between alleles in the alternative sex chromosomes. Such evolutionary strata have long been thought to result from sexually antagonistic selection, i.e., the successive recruitment of genes with alleles beneficial only in one sex but harmful in the other, where selection favors complete linkage to the sex-determining genes (Charlesworth, 2017). However, little evidence could be found in support of this hypothesis (Ironside, 2010) and recent studies have demonstrated the existence of recombination suppression across large regions and of multiple evolutionary strata on fungal mating-type chromosomes despite the lack of such antagonistic selection (Bazzicalupo et al., 2019; Foulongne-Oriol et al., 2021; Hartmann et al., 2021; Menkis et al., 2008). Fungal mating-type chromosomes thus represent valuable emerging models for understanding how and why recombination suppression evolves (Hartmann et al., 2021).

Anther-smut fungi in the *Microbotryum violaceum* species complex (Basidiomycota) are particularly good models to address these questions. These plant-castrating fungi have an obligate sexual cycle of meiosis and mating prior to infecting each new host, and they have megabase-long non-recombining regions on their mating-type chromosomes (Badouin et al., 2015; Branco et al., 2018, 2017; Carpentier et al., 2019; Hartmann et al., 2019; Hood et al., 2004). This recombination suppression evolved independently at different times across the *Microbotryum* phylogeny, providing independent cases of evolution following recombination suppression with various ages (0.20-2.30 MY; Branco et al., 2018, 2017). Furthermore, in multiple *Microbotryum* species, recombination suppression has extended outward from the mating-type genes in several successive steps (Branco et al., 2018, 2017).

Most basidiomycete fungi carry two mating-type loci on different chromosomes and mating can only be successful between gametes harboring different alleles at both loci. The PR mating-type locus includes pheromone and pheromone receptor genes controlling gamete fusion. The HD mating-type locus has two homeodomain genes controlling post-mating growth. In *Microbotryum* anther-smut fungi, the PR and HD loci were ancestrally located on different chromosomes (Branco et al., 2017). The different genes within each locus have been linked since the emergence of the basidiomycete clade (Coelho et al., 2017; Sun et al., 2019), and recombination suppression extended in the local region around each of the two mating-type loci early in the diversification of *Microbotryum* fungi, generating independent evolutionary strata around the PR locus (called purple stratum) and HD locus (called blue stratum), and later another stratum (called orange) distal to the purple stratum (Branco et al., 2017). Subsequently, at least eight independent events of recombination suppression involving different chromosomal rearrangements linked the HD and PR loci (Branco et al., 2018; Carpentier et al., 2021). The linkage of PR to HD mating-type determining genes is beneficial under the highly selfing system of *Microbotryum* fungi; only two mating types are then produced by a diploid individual, leading to a 1/2 compatibility among gametes under inter-tetrad selfing, i.e. among gametes produced by different meiosis events of a single diploid individual, and 2/3 compatibility under intra-tetrad selfing, i.e. among products of the very same meiosis. The linkage of the HD and PR mating-type loci thus doubles the odds of gamete compatibility under inter-tetrad selfing compared to a system with unlinked loci, in which a diploid individual produces four mating types, i.e. with 1/4 compatibility among gametes (Branco et al. 2017). In two *Microbotryum* fungi, the PR and HD loci became linked, not to each other, but to their respective centromeres; while this is not beneficial under inter-tetrad selfing (1/4 compatibility), it yields the same 2/3 compatibility odds under intra-tetrad selfing as PR-HD linkage (Carpentier et al., 2019). In previous studies, we used the term “black strata” in referring all genomic regions linking the PR to HD mating-type loci to each other or to respective centromeres, although it should be reminded that they correspond to independent evolutionary events, trapping different sets of genes, with subsets overlapping among some of the black strata. HD-PR linkage was further followed, in several species, by stepwise extension of recombination suppression beyond the mating-type loci. The evolutionary and proximal causes for such extensions are still unknown (Hartmann et al., 2021); a recent theoretical model suggests that recombination suppression beyond mating-type loci may evolve due to the benefit of sheltering deleterious mutations (Jay et al., 2021). The young evolutionary strata generated by these recombination suppression extensions in *Microbotryum* fungi were called by different colors in previous studies, e.g., white, light blue, pink, green and red (Branco et al., 2018).

The alternate alleles at genes in the non-recombining regions of mating-type chromosomes independently accumulate substitutions with time; their divergence is thus a proxy of time since the cessation of recombination. For detecting distinct evolutionary strata, one typically uses non-synonymous substitutions (d_S_), i.e. not changing the amino-acid sequence, as they are considered neutral. In selfing species such as *Microbotryum*, being highly homozygous, the synonymous divergence between two alleles in recombining regions is null or very close to zero. The d_S_ can therefore be used to delimit the non-recombining regions (NRRs) and the recombining pseudo-autosomal regions (PARs) of the mating-type chromosomes. Trans-specific polymorphism is also used to study the age of recombination suppression linking genes to mating-type loci. Indeed, as soon as a gene is fully linked to a mating-type locus, its alternative alleles will remain associated to alternative mating types (called a_1_ and a_2_ in *Microbotryum* fungi), even across speciation events; in a genealogy, its alleles will therefore be grouped according to mating type rather than according to species, and the nodes at which the alleles associated with the alternative mating types diverge indicate the time of recombination cessation (Branco et al., 2017, 2018; Hartmann et al., 2021).

In sex-determining homogametic/heterogametic systems, one chromosome still recombines, thus used as a proxy for the ancestral gene order (Lahn and Page, 1999). In contrast, because mating types are determined at the haploid stage in fungi, both mating-type chromosomes are always heterozygous and both undergo frequent rearrangements in non-recombining regions, which renders the inference of historical steps of recombination suppression challenging. Previous studies have used as proxy of the ancestral state the chromosomal arrangement and gene order of an outgroup, *M. intermedium*, carrying its PR and HD loci on different chromosomes and having its gene order highly syntenic with two other distantly related species, *M. lagerheimii* and *M. saponariae*, also carrying their PR and HD loci on different chromosomes (Branco et al., 2017; Carpentier et al. 2019). However, the genome of a single mating type of *M. intermedium* was available so far.

In this study, we therefore sequenced the genome of the lacking mating type of *M. intermedium*, as well as the haploid genomes of the two mating types of four additional species displaying 2/3 compatibility between gametes within tetrads, suggesting either the linkage of the PR to the HD mating-type loci or the linkage of each mating-type locus to their respective centromeres (M.E. Hood, unpublished data). Among these four species, *M. v. viscidula, M. v. gracilicaulis* and *M. v. parryi* belong to clades with *Microbotryum* fungi carrying linked HD and PR mating-type loci, while *M. v. lateriflora* belongs to the clade with the two species with HD and PR loci on distinct chromosomes, i.e., *M. lagerheimii* and *M. saponariae*. We analysed the organisation of the mating-type chromosomes in the new genomes sequenced in order to address the following questions: 1) Does *M. v. lateriflora* also carry its PR and HD mating-type loci on distinct chromosomes, with or without recombination suppression with the respective centromeres, or does it correspond to yet another independent event of HD-PR linkage? 2) Do the three other species with new genome sequenced display linked HD and PR loci, and do the phylogenetic placements and trans-specific polymorphism indicate additional HD-PR linkage independent events from the previously documented ones? 3) In case of independent events of HD-PR linkage, did they occur through the same chromosomal rearrangements or yet new types of rearrangements? 4) Are there footprints of evolutionary strata in the new genomes analysed, i.e., stepwise extension of recombination suppression?

## Material and methods

### DNA extraction and sequencing

DNA extraction and sequencing based on PacBio (Pacific Bioscience) long-read sequencing was performed as described previously (Branco et al., 2018, 2017). Samples were collected before the publication of laws regarding the Nagoya protocol, if any, in the countries of collection. *Microbotryum violaceum* is a species complex, with most species being specialized on a single host species, especially when parasitizing *Silene* plant species (Hartmann et al., 2019; Le Gac et al., 2007a, 2007b). For the species of the complex that have not been formally named yet, they are named after the species name of the host plant, as commonly practiced for host races or *formae speciales* in phytopathology. For the present study, a_1_ and a_2_ haploid cells were isolated from a single tetrad from the following species: *M. violaceum lateriflora* parasitizing *Moehringia lateriflora* (strain 1509, Boothbay Harbor, Maine, USA, Coord.: 43°51’34.8”N 69°37’45.6”W, collected in 2017), *M. violaceum gracilicaulis* parasitizing *Silene gracilicaulis* (strain 1299, Wutoudi, Lijiang, China, Coord: 27°03’01.9”N 100°11’36.5”E, collected in 2014), *M. violaceum parryi* parasitizing *S. parryi* (strain 1510, Waterton lakes, Canada, Coord.: 49°1’48N 113°50’24”W, collected in 2017), and *M. violaceum viscidula* parasitizing *S. viscidula* (strain 1506, Ninglang, Lijiang, China, Coord.: 27°12’18.7” 100°47’35.7”, collected in 2015). Cultivation of a_1_ and a_2_ haploid cells was performed as in Le Gac al., 2007b. We sequenced the haploid genome of *M. intermedium* corresponding to the a_2_ mating type of the very same strain for which the haploid genome of the a_1_ mating type had already been sequenced (strain 1389-BM-12-12, collected on *Salvia pratensis*, Italy, Coord. GPS : 44°20’00.7”N 7°08’10.9”E; Branco et al. 2017).

### Genome assemblies and gene prediction

Raw reads were processed using tools from the smrtanalysis suite 2.3.0 (https://github.com/PacificBiosciences/GenomicConsensus) as the previously published genomes (Badouin et al., 2015; Branco et al., 2018, 2017). We converted the bax.h5 files from the same sequencing run into one fastq file using pls2fasta. We generated the assembly using canu v.1.8 (Koren et al., 2017) with the parameters “genomeSize=30m” and “-pacbio-raw”. We used pbalign (version 0.3.0) with the blasr algorithm (Chaisson and Tesler, 2012) to realign the raw reads onto the assembly (indexed with samtools faidx; Li et al., 2009) and then used the output bam file into quiver (Chin et al., 2013) to polish the assembly basecalling. Default parameters were used when not detailed in the text. See Supplementary Table S1 for assembly statistics.

As for the previously published *Microbotryum* high-quality genome assemblies (Badouin et al., 2015; Branco et al., 2018, 2017; Carpentier et al., 2019), the protein-coding gene models were predicted with EuGene 4.2a (Foissac et al., 2008), trained for *Microbotryum*. Similarities to the fungal subset of the uniprot database (Consortium TU, 2011) plus the *M. lychnidis-dioicae* Lamole proteome (Badouin et al., 2015) were integrated into EuGene for the prediction of gene models.

### Transposable element detection and annotation

D*e novo* detection of transposable elements (TEs) was done using LTRharvest (Ellinghaus et al., 2008) from GenomeTools 1.5.10, performing long-terminal repeat (LTR) retrotransposons detection and RepeatModeler 1.0.11 (Smit and Hubley, 2008) combining results from three other programs, RECON (Bao and Eddy, 2002), RepeatScout (Price et al., 2005) and Tandem Repeats Finder (Benson, 1999). The TE detection was enriched by BLASTn 2.6.0+ (Altschul et al., 1990) using the genomes as a database and the previously detected TE models as queries. To fulfill the repetitivity criterion, a TE sequence detected by RepeatModeler or LTRHarvest and their BLAST hits were retained only if the query matched three or more sequences in the same species with an identity ≥ 0.8, a sequence length > 100bp and a coverage (defined as the query alignment length with removed gaps divided by the query length) ≥ 0.8. When these criteria were met, the other query matches were retained for the following parameters: identity ≥ 0.8, sequence length > 100bp, e-value ≤ 5.3e-33 and coverage ≥ 0.8. TE annotation (Wicker et al., 2007) was performed using the fungal Repbase database 23.05 (Bao et al., 2015) based on sequence similarity. First, a BLASTn search and a BLASTx search were performed using the fungal Repbase DNA and protein sequence databases, respectively, and TE sequences as queries. For each of these similarity-based searches, the minimum e-value score was set at 1e−10 and minimum identity at 0.8. Then, protein domain detection was performed using pfam_scan.pl (ftp://ftp.ebi.ac.uk/pub/databases/Pfam/Tools/) on TE sequences and compared to protein domain detection in the fungal Repbase database. Matches with e-values lower or equal to 1e-5 were kept. The annotation found by RepeatModeler was also considered. When no annotation was found, the TE sequence was annotated as “*unclassified*”; such annotations were discarded when overlapping with predicted genes. Annotation was performed using a python script. TE copies were removed from the predicted genes in further analyses.

### Comparative genomics

Orthologous groups were obtained by Markov clustering (van Dongen, 2000) of high-scoring pairs parsed with orthAgogue (Ekseth et al., 2014) based on all-vs-all BLASTP 2.6.0 (Altschul et al., 1990) on protein sequences. The coding sequences (CDS) of 3,645 single-copy genes present in all species analysed were aligned independently using MUSCLE (Edgar, 2004) as implemented in TranslatorX v1.1 (Abascal et al., 2010). We used IQ-TREE 2.0.4 (Minh et al., 2020) to build the maximum likelihood species tree with the *TIM3+F* model of substitution chosen according to the Akaike information criterion (AIC) by Model Finder implemented in IQ-TREE (Kalyaanamoorthy et al., 2017). We used *Rhodothorula babjavae* as an outgroup. We assessed the robustness of the nodes with 1000 ultrafast bootstraps (Hoang et al., 2018; Minh et al., 2013) and SH-like approximate likelihood ratio tests (SH-aLRT) (Guindon et al., 2010) from the concatenated alignment. Pheromone-receptor (PR), pheromone and homeodomain (HD) genes were identified by BLASTN 2.6.0 (Altschul et al., 1990) similarity search using the pheromone receptor, pheromone and homeodomain gene sequences from *M. lychnidis-dioicae* (Devier et al., 2009; Petit et al., 2012). We identified the contigs belonging to the mating-type chromosomes for each species by i) identifying the contigs carrying PR, pheromone and HD genes, ii) identifying the contigs carrying single-copy orthologous genes from the PR and HD chromosomes of *M. intermedium*, iii) identifying the contigs in the a_1_ (respectively a_2_) genome carrying single-copy orthologous genes homologous to genes on the identified mating-type contigs of the a_2_ (respectively a_1_) genome, iv) plotting all these contigs using circos (http://circos.ca) comparing a_1_ and a_2_ genome to check which contigs really belong to the mating-type chromosomes (the contigs fully collinear between mating types and not assembled at an edge of the mating-type chromosome were considered to belong to autosomal chromosomes, see Branco et al 2017). We oriented contigs in the direction involving the fewest inversions and keeping the PARs at the edges and non-inverted between mating-type chromosomes within a diploid genome. The contigs were considered joined when not broken at the same point in the alternative mating-type genomes within diploid individuals. Mating-type fission and fusion scenarios were inferred by linking orthologous genes on a circos plot between the mating-type contigs from one haploid genome with linked mating-type loci and the mating-type chromosomes of *M. intermedium* or *M. lagerheimii*, used as a proxy for the ancestral state as done previously (Branco et al., 2018, 2017).

Synonymous divergence (d_S_) was calculated from the alignment of a_1_ and a_2_ allele sequences using MUSCLE (Edgar, 2004) implemented in TranslatorX v1.1 (Abascal et al., 2010). Synonymous divergence (d_S_) and its standard error were calculated using the yn00 v4.9f program of the PAML package (Yang, 2007) and plotted using the ggplot2 library of R (Wickham, 2009). Non-recombining regions were identified as non-collinear regions with non-null synonymous divergence using the a_1_-vs-a_2_ circos and d_S_ plots. Pseudo-autosomal regions were identified as the remaining regions, being collinear and with null synonymous divergence, as expected in these highly selfing fungi.

Centromeres (Supplementary Table S2) were identified by blasting (BLASTN 2.6.0 Altschul et al., 1990) the centromeric repeats previously described (Badouin et al., 2015). Centromeres were defined as stretches of centromeric repeats (Supplementary Table S3), two consecutive repeats being distant by maximum 50kb; only the largest block was kept when several stretches of centromeric repeats were found on the same contig. Only contigs larger than 20kb were considered. The delimitation of putative centromeres was recursively extended by 1kb windows as long as gene density was lower than 0.25 in the focal window. Results were congruent with previously identified centromeres (Branco et al., 2018). Telomeres (Supplementary Table S4) were delimited in the 100bp region at the end of contigs carrying at least five times the telomere-specific TTAGGG motif (CCCTAA on reverse complementary strand) as done previously (Badouin et al., 2015).

### Trans-specific polymorphism and datation

We performed codon-based alignment with macse v2.0 (Ranwez et al., 2018) of one-to-one orthologs of genes ancestrally located between the PR locus and its centromere (23 genes), between this centromere and the short arm telomere (11 genes), between the HD locus and its centromere (8 genes) or in the light blue *M. caroliniana* stratum (14 genes). Gene trees were obtained with IQ-TREE 2.0.4 (Minh et al., 2020) with automatic selection of the substitution model (Kalyaanamoorthy et al., 2017) and branch-support estimated with 1000 ultrafast bootstraps (Hoang et al., 2018; Minh et al., 2013). Tree topology congruence was assessed with the approximate unbiased (AU) test (Shimodaira, 2002) with 10,000 RELL replicates (Kishino et al., 1990) implemented in IQ-TREE 2.0.4 (-zb 10000 and -au options). For the genes ancestrally located between the PR locus and its centromere, between the centromere of the ancestral PR chromosome and its short arm telomere and between the HD locus and its centromere we tested whether each gene alignment was significantly conflicting with each of the different topologies obtained among the 41 other gene trees, using the -z option and the substitution model fixed to the one found during the focal gene tree reconstruction. We also compared the topology of the species tree and the tree obtained by concatenating the conserved genes in the region proximal to the PR locus corresponding to light blue and red strata. Trees with pAU values greater than 0.05 were considered non-discordant. The codon-based multiple alignments were concatenated separately for these regions, producing alignments of 19,944 (PR locus to centromere), 3,955 (centromere ancestral PR to short-arm telomere), 7,705 (HD locus to centromere) and 9,952 (proximal region of the PR locus on the long arm side, corresponding to the light blue and red strata) codons. Partitionated alignments (one partition per codon position) were imported to BEAUti v2.5.0 (Bouckaert et al., 2014) to produce BEAST2 xml input files with the following priors: unlinked gamma site model with 3 categories and HKY substitution model; strict clock and gamma clock rate with alpha 0.001 and beta 1,000; calibrated Yule tree model with gamma birth alpha 0.001 and beta 1,000; exponential gamma shape per partition; log-normal HKY’s kappa. A calibration point at each of the two divergence nodes between *M. lychnidis-dioicae* and *M. silenes-dioicae*, one node separating the a_1_-associated alleles and the other node the a_2_-associated alleles, both set as normally distributed with mean 0.42 MY and sigma 0.04. BEAST2 v2.4.6 (Bouckaert et al., 2014); runs were performed for 20,000,000 iterations and written every 1,000 iterations. Date estimates were obtained with the 15,000 trees with best posterior probabilities.

## Results

We obtained high-quality assemblies (Supplementary Table S1) for a genome of the previously unavailable haploid mating type (a_2_) of *M. intermedium*, as well as the haploid genomes of both mating types of four additional species compared to earlier studies: *M. violaceum lateriflora* parasitizing *Moehringia lateriflora, M. violaceum gracilicaulis* parasitizing *Silene gracilicaulis, M. violaceum parryi* parasitizing *S. parryi*, and *M. violaceum viscidula* parasitizing *S. viscidula*. The species tree obtained based on 3,645 single-copy genes was robust and congruent with previous studies; the new species were distributed across the phylogeny (Figure 1).

**Figure 1:**
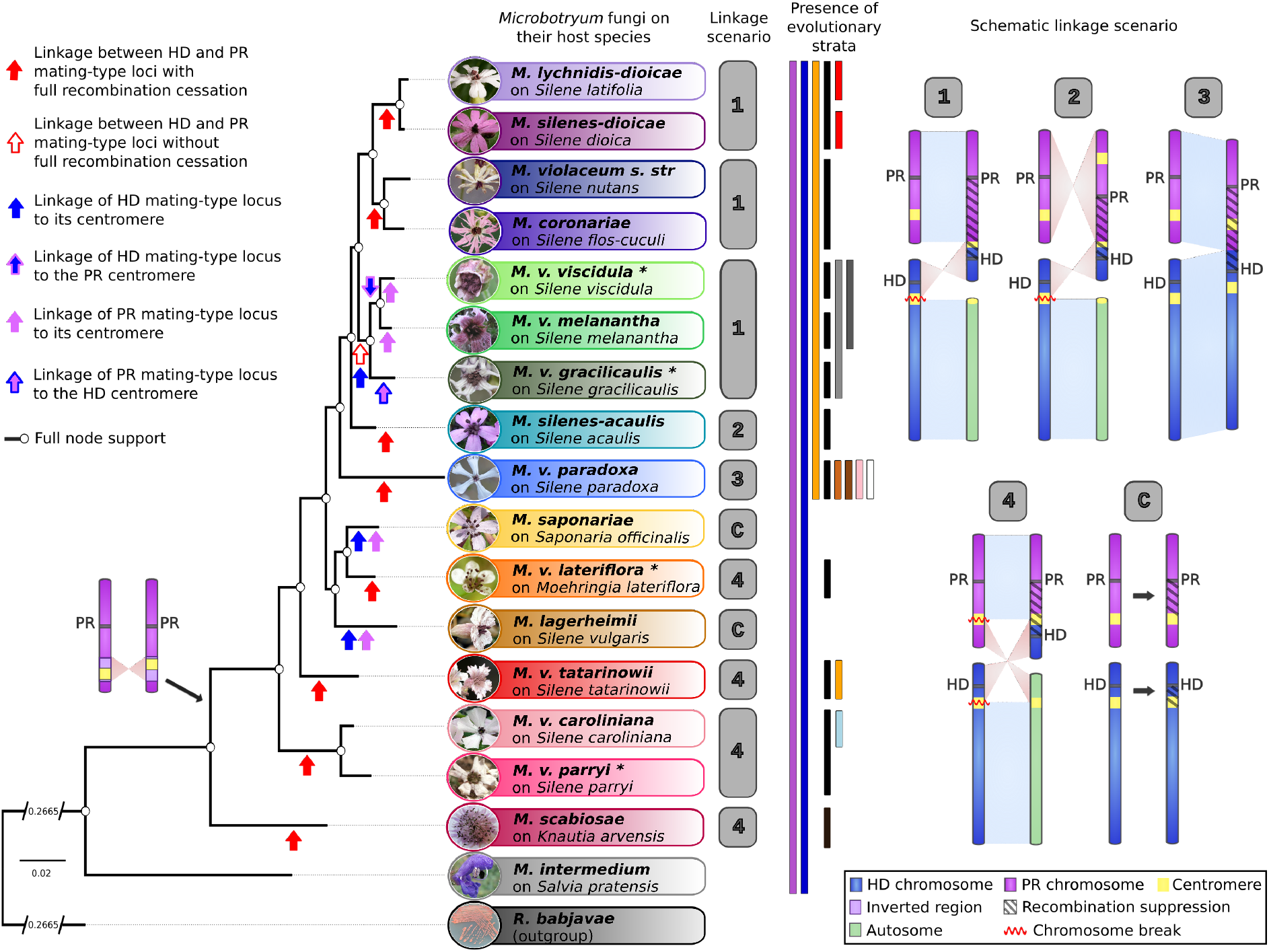
Multiple independent events of recombination suppression in mating-type chromosomes across the *Microbotryum* phylogeny. Phylogeny of 17 *Microbotryum* species rooted by a red yeast outgroup (*Rhodothorula babjavae*), based on single-copy orthologous gene genealogies (left panel), with pictures of diseased plants. White circles at nodes indicate full support by bootstrap and Shimodaira–Hasegawa-approximate likelihood ratio tests (SH-aLRT). The genomes of the species labelled with * were generated for this study. Photos of the diseased flowers by M. E. Hood, except *M. v. viscidula* and *M. v. gracilicaulis* (by H. Tang) and *M. v. parryi* (by acorn13 @iNaturalist, cropped). CC BY 4.0. The red arrows represent the independent events of pheromone-receptor (PR) and homeodomain (HD) mating-type locus linkage. The blue and purple arrows represent the independent events of HD and PR mating-type locus linkage to their respective centromeres, respectively. The purple-framed blue arrow represents the event of HD mating-type locus linkage to the ancestral PR centromere and the blue-framed purple arrow represents the events of PR mating-type locus linkage to the ancestral HD centromere. The chromosomal fission and fusion scenarios having led to HD-PR linkage are indicated right to the species tree (labeled with numbers, in grey squares corresponding to independent events). The scenario labeled “C” corresponds to recombination suppression events of mating-type loci to their respective centromeres and is depicted on the right panel. The color bars correspond to the presence of the different evolutionary strata in each species. On the depicted chromosomes are represented in purple the ancestral or fused part of the PR chromosome, in blue the ancestral or fused part of the HD chromosome, in green the regions of the ancestral PR and HD chromosomes that became autosomal following HD-PR linkage. The inversion having occurred near the centromere of the PR chromosome is represented in light purple. The centromeres are represented in yellow. Non-recombining regions (NRRs) are cross-hatched. Chromosome fissions are indicated by red waves.

### Ancestral state in Microbotryum *intermedium*

The assembly of the a_2_ *M. intermedium* genome recovered full HD and PR mating-type chromosomes, with telomeric repeats at the ends of the contigs and well-defined centromeres, thus confirming that the PR and HD loci are unlinked in this species. We found, as in previous studies on other *Microbotryum* species (Branco et al. 2017, 2018), that autosomes were completely collinear between the two haploid genomes of the sequenced *M. intermedium* diploid strain (Supplementary Figure S1A), and autosomal genes had zero d_S_ levels between the allele copies (Supplementary Figure S2A); d_S_ levels and collinearity are therefore good indicators of the occurrence of recombination and these selfing species are highly homozygous. By comparing the alternative mating-type chromosomes of *M. intermedium*, we detected few rearrangements (Figure 2A) and the d_S_ plots mostly displayed zero values, as observed in autosomes (Supplementary Figures S1A and S2A). These observations confirm that there is likely no recombination suppression along most of the *M. intermedium* mating-type chromosomes. High d_S_ values and rearrangements were only observed in proximity to the PR locus, indicating that this region does not recombine, corresponding to the stratum called “purple” that was previously identified in the other *Microbotryum* species (Figure 3A). This purple evolutionary stratum thus likely evolved before the divergence of *M. intermedium* from the other *Microbotryum* species (Figure 1).

**Figure 2:**
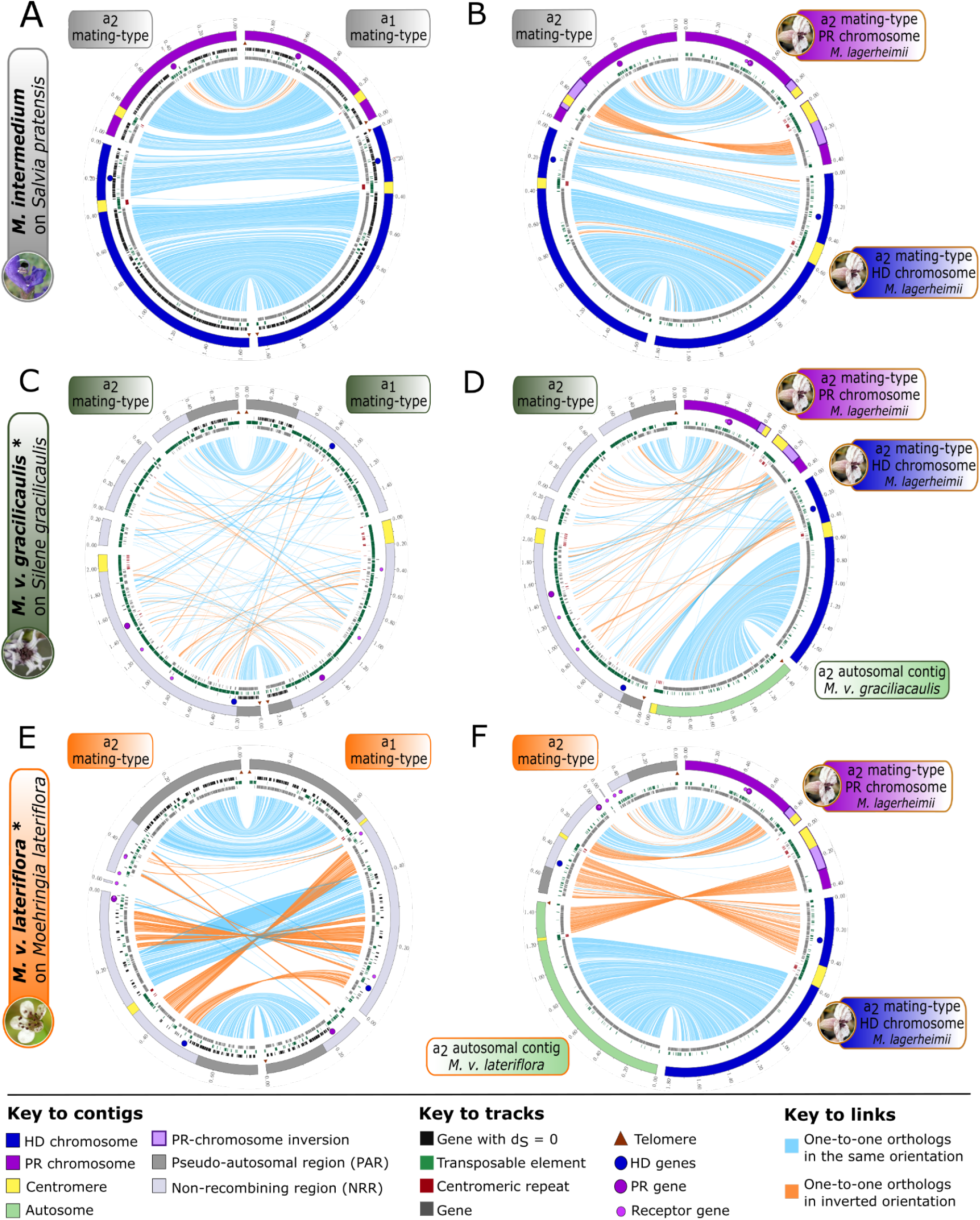
Inter- and intra-specific comparison of gene order between mating-type chromosomes in *Microbotryum* fungi. Synteny plots between A) homeodomain (HD) and B) pheromone-receptor (PR) chromosomes of opposite mating types in *M. lagerheimii*, C) a_1_ and a_2_ mating-type chromosomes of *M. v. gracilicaulis*, D) a_2_ mating-type chromosome of *M. lagerheimii* and their homologues in *M. v. gracilicaulis*, E) a_1_ and a_2_ mating-type chromosomes of *M. v. lateriflora*, F) a_2_ mating-type chromosomes in *M. lagerheimii* and their homologues in *M. v. lateriflora*. Comparisons of *M. lagerheimii* a_2_ HD and PR chromosomes and *M. v. gracilicaulis* (D) and *M. v. lateriflora* (F) a_2_ mating-type contigs to infer chromosomal rearrangements having led to HD-PR linkage, considering *M. lagerheimii* chromosomes as a proxy for the ancestral chromosomal state. The PR, HD and pheromone genes are represented by purple, blue and small light purple circles, respectively. The outer track represents contigs, with length ticks every 200 kb. The HD and PR mating-type chromosomes of *M. lagerheimii* are represented in blue and purple, respectively (A, B, D, F). The region of the inversion having occurred after divergence with *M. scabiosae* (Fig. 1), encompassing the centromere of the PR chromosome, is highlighted in light purple. For *M. v. gracilicaulis* (C,D) and *M. v. lateriflora* (E,F), the non-recombining region (NRR) and the pseudo-autosomal region (PAR) of the mating-type chromosomes are represented in light and dark gray, respectively. Green contigs correspond to autosomal contigs of *M. v. gracilicaulis* (D) and *M. v. lateriflora* (F). The centromeres are represented in yellow. The telomeres are indicated by brown triangles. Black marks on the inner track represent the genes with null synonymous divergence between a_1_ and a_2_ alleles. Green marks on the inner track represent transposable element (TE) copies. Gray marks on the inner track correspond to genes. Red marks on the inner track correspond to the putative centromeric repeats. Blue and orange lines link alleles, the latter corresponding to inversions. The link width is proportional to the corresponding gene length.

**Figure 3:**
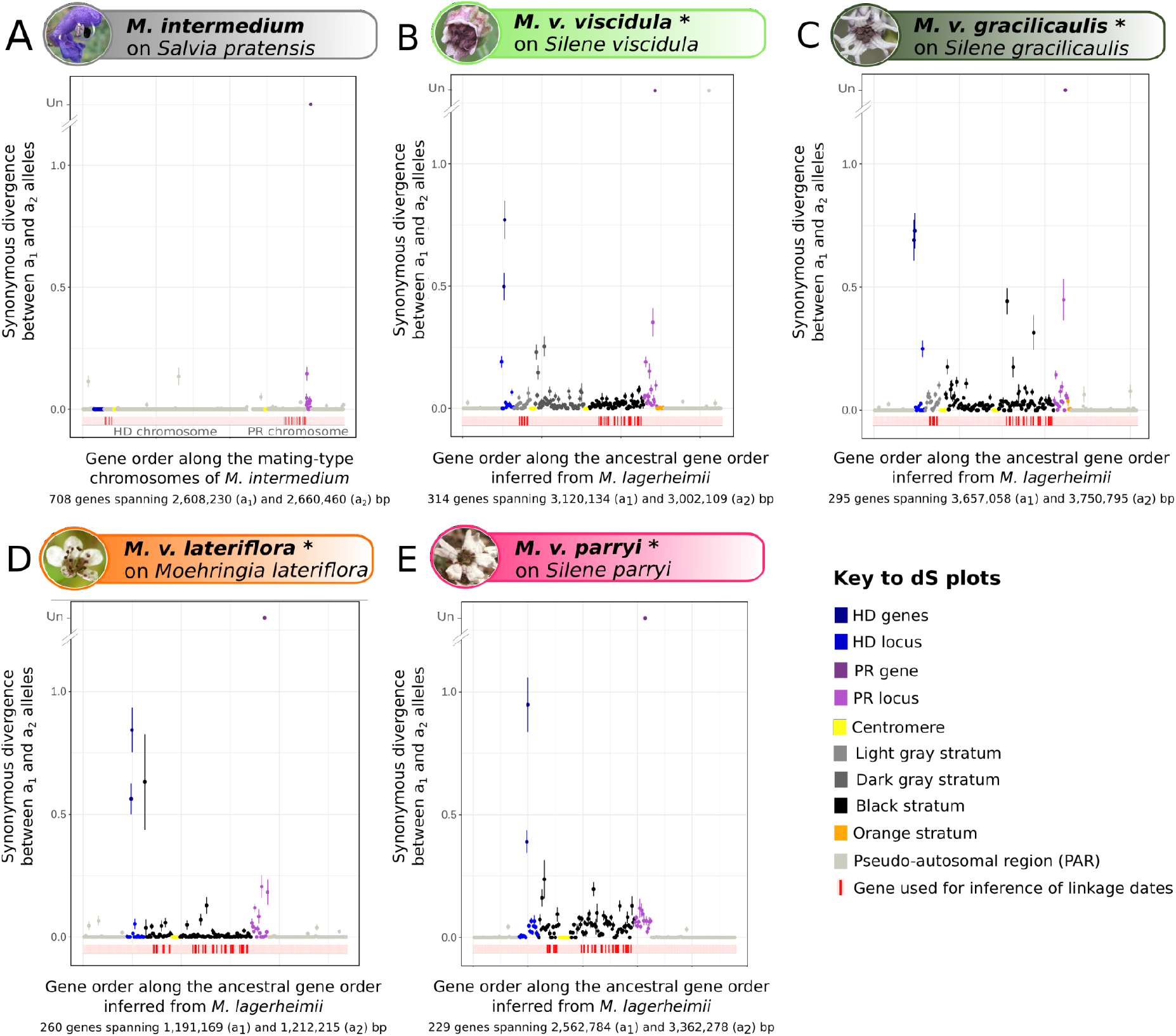
Evolutionary strata in the mating-type chromosomes of *Microbotryum* fungi. Per-gene synonymous divergence between a_1_ and a_2_ alleles on the mating-type chromosomes of A) *M. intermedium*, B) *M. v. viscidula*, C) *M. v. gracilicaulis*, D) *M. v. lateriflora* and E) *M. v. parryi*, plotted along the ancestral gene order, inferred from *M. lagerheimii* with unlinked mating-type loci. The number of genes and average of the a_1_ and a_2_ contig lengths are indicated on the X axis. Synonymous divergence is used as a proxy for time since recombination suppression. Ancient purple and blue evolutionary strata were formed around each of the mating-type genes (pheromone-receptor, PR, in dark purple, and homeodomain, HD, in dark blue, genes controlling mating compatibility) and is ancestral in the *Microbotryum* clade (present in all species, including *M. intermedium* in A). In *M. v. viscidula* (B) recombination suppression first linked the HD locus to its centromere, then extended to the PR centromere and eventually to the PR locus, generating the light gray, dark gray and the black strata. In *M. v. gracilicaulis* (C), recombination suppression linked the HD locus to its centromere and then to the PR locus, forming the light gray and the black strata. In *M. v. lateriflora* (D) and *M. v. parryi* (E), recombination suppression has linked the HD and PR mating-type loci together, generating black strata. Further extension of recombination suppression beyond the mating-type loci generated the younger orange evolutionary stratum in *M. v. viscidula* (B) and *M. v. gracilicaulis* (C). The recombining regions of the mating-type chromosomes with null synonymous divergence, called pseudo-autosomal regions (PARs), are represented in gray. Red ticks below the X axis correspond to genes used to infer the linkage dates.

We found a small inversion near the centromere in the PR mating-type chromosome that distinguished *M. intermedium* from another species also with PR and HD loci retained on separate mating-type chromosomes, *M. lagerheimii* (Figure 2B). Because the *M. lagerheimii* arrangement was found in all the *Microbotryum* species analysed except *M. intermedium* and *M. scabiosae* (Figure 2B and Supplementary Figure S3O), we inferred that *M. intermedium* harbored the ancestral gene order and that the inversion around the PR centromere occurred early in the *Microbotryum* clade, after the divergence of the rest of the *Microbotryum* clade from the *M. scabiosae* lineage (Figure 1). We therefore consider the *M. lagerheimii* gene order as ancestral for inferring chromosomal rearrangement scenarios in the lineages that have derived subsequent to this inversion near the PR chromosome centromere (Figure 1).

### Discovery of additional independent events of mating-type loci linkage

The evolutionary history of mating-type chromosomes was inferred in four *Microbotryum* species studied for the first time here (Figure 1, *M. v. gracilicaulis, M. v. viscidula, M. v. lateriflora* and *M. v. parryi*) by comparing their genomes to the *M. lagerheimii* genome. We obtained high-quality assemblies of two haploid genomes of opposite mating types per species, i.e. haploid genomes of alternative mating types (a_1_ and a_2_) produced by meiosis of a single diploid individual. The mating-type chromosomes were assembled in few contigs, not interrupted at the same places in the two mating-type chromosomes in each species (Figure 2 and Supplementary Figure S4), which allowed complete joining of contigs. The PR and the HD loci were found on a single contig in at least one haploid genome of each species, indicating that these species all have their mating-type loci linked together on a single chromosome (Figures 2C and 2E; Supplementary Figures S4A and S4C). The pseudo-autosomal regions were defined as fully collinear regions lacking synonymous divergence between alleles (i.e. with the same patterns as in autosomes; Figure 2; Supplementary Figures 1 and 2).

We reconstructed the evolutionary history of their mating-type chromosomes by comparing their genome structures to those of *M. lagerheimii*, taken as a proxy for their genomic ancestral state before chromosomal fusion events. The mating-type chromosome fissions (always found at centromeres) were determined by assessing, in synteny plots (e.g. Figure 2 and Supplementary Figure S4), what arms of the ancestral mating type chromosomes became autosomes, i.e. were completely collinear between mating-types and assembled separately from the derived mating-type chromosomes in both haploid genomes. The orientation of ancestral mating-type chromosome fusion was assessed by determining on synteny plots (e.g. Figure 2 and Supplementary Figure S4) what edge of ancestral mating-type chromosomes remained recombining, i.e. became PARs, the other edge of the ancestral chromosome or the centromere thus corresponding to the fusion point. For orienting the few contigs within the non-recombining regions that could not be assembled with the PAR contigs, we applied a majority rule minimizing inversion numbers with the alternative mating type; their orientations do not impact the scenario reconstruction.

In *M. v. gracilicaulis* parasitizing *Silene gracilicaulis* (Figure 2D) and *M. v. viscidula* parasitizing *Silene viscidula* (Supplementary Figure S3B), mating-type locus linkage was achieved through the fusion of the entire PR chromosome and the short arm of the HD chromosome. This represents the same chromosomal rearrangement as in *M. lychnidis-dioicae, M. silenes-dioicae, M. coronariae, M. violaceum s*.*s*. and *M. v. melanantha*. The placement of these species as a clade in the phylogeny suggests that this fission/fusion event can represent an ancestral rearrangement to this clade (Figure 1). Under the alternative hypothesis, the same rearrangement would have occurred several times independently.

In order to test the independence of the mating-type locus linkage events in this clade, we analysed trans-specific polymorphism by reconstructing the evolutionary history of a_1_ and a_2_-associated alleles for the genes located between the two mating-type loci at the time of the linkage event. We built the genealogies of 23, 11 and 8 genes ancestrally located between the PR locus and its centromere, in the PR-chromosome short arm, and between the HD locus and its centromere, respectively. Among these 42 genes, the 31 (23+8) ancestrally in the PR-to-centromere and HD-to-centromere regions were initially located between the two mating-type loci for all types of rearrangements documented so far, except in *M. silenes-acaulis*, in which the PR chromosome was fused in the reverse orientation so that the genes ancestrally located between the PR locus and its centromere are still recombining. The genealogies recovered in the genes ancestrally located between the PR locus and its centromere and between the HD locus and its centromere displayed trans-specific polymorphism between *M. lychnidis-dioicae* and *M. silenes-dioicae* on the one hand and *M. violaceum s. str*. and *M. coronariae* on the other hand. This indicates that recombination cessation occurred independently in these two clades (Figure 1), either following independent similar chromosomal rearrangements or a single basal chromosomal rearrangement with initially incomplete recombination suppression. We also found trans-specific polymorphism, shared between *M. v. caroliniana* and *M. v. parryi*, in the HD-to-centromere and PR-to-centromere regions, suggesting: 1) an ancestral chromosomal rearrangement and gradual completion of recombination suppression, or 2) recombination suppression between each mating-type locus and its respective centromere, as in *M. lagerheimii* and *M. saponariae*, and then, indepently in the two species, the same type of chromosomal rearrangement linking the HD and PR loci one two each other. The two steps would be beneficial, as the linkage between each mating-type locus and its respective centromere is beneficial under intra-tetrad selfing, while the HD-PR linkage is, in addition, also beneficial under inter-tetrad selfing (Carpentier et al. 2019). Both alternatives are consistent with much older date estimates for recombination suppression in both regions (2.23 or 2.79 MYA) than the inferred speciation date (1.42 MYA, Figure 4 and Supplementary table S5).

**Figure 4:**
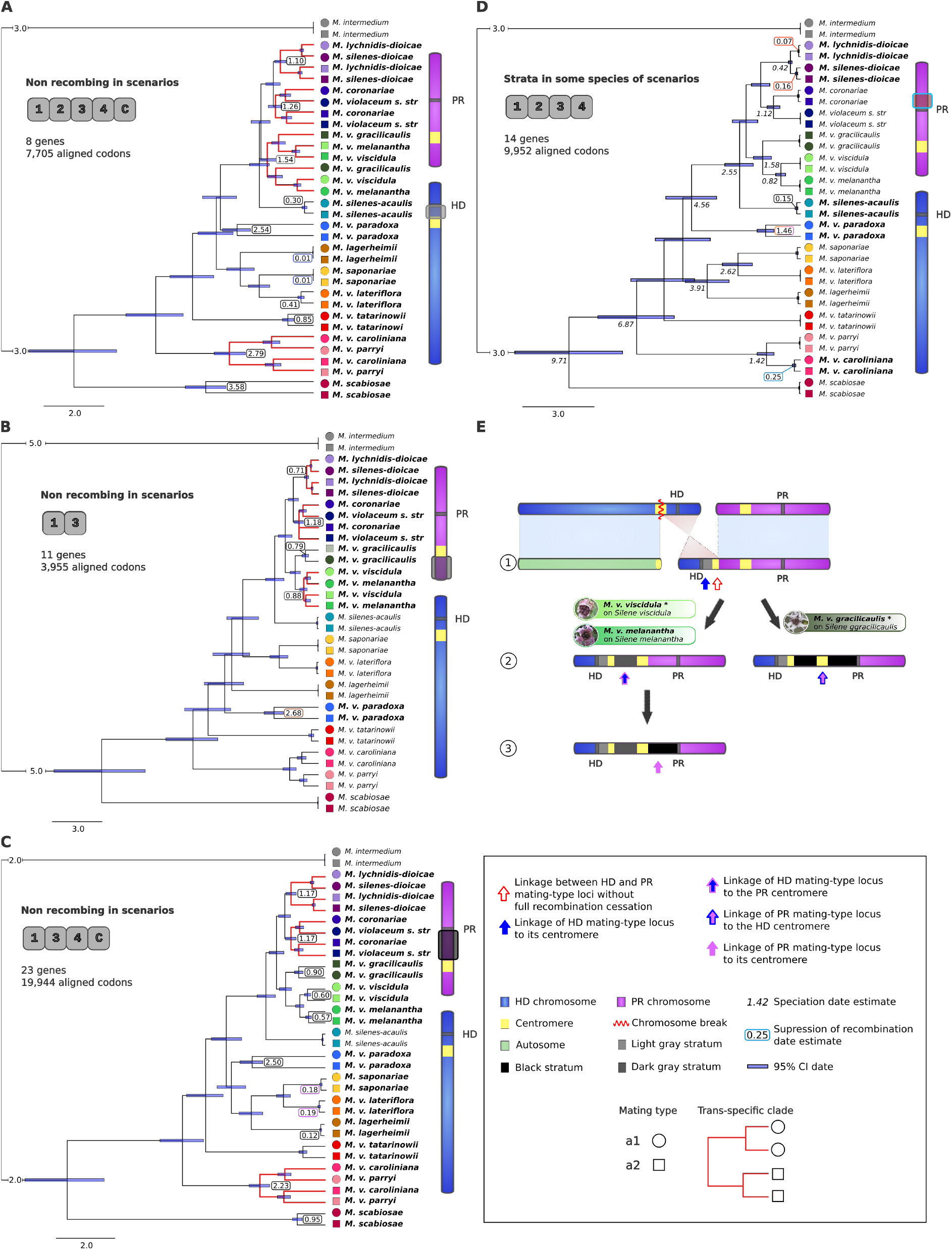
Onset of suppression of recombination date estimates and reconstructed scenario of stepwise recombination suppression between the HD and PR mating-type loci in the *Microbotryum violaceum melananth*a clade. A-D) Timetree of conserved genes in the regions boxed on the right-side ancestral chromosome sketches. Topologies are all significantly different (AU test p-value > 0.05). Trans-specific polymorphisms are indicated by red branches. Date estimates are shown in colored boxes near the split between a_1_ and a_2_ alleles, confidence intervals (CI) depicted as rectangles at corresponding nodes. Numbers in italics below branching points in D correspond to speciation dates. Topology in D is identical to the species phylogeny and estimated speciation dates overlap. Insets detail the scenarios in which the conserved genes are in non-recombining regions (species names in bold), the number of genes concatenated and the alignment length in codons. See Figure 1 for diagrams of the reconstructed scenarios. Colored symbols correspond to a_1_ (circles) and a_2_ alleles (squares) following the species color scheme in Figure 1. E) Reconstructed stepwise recombination suppression having generated the light gray, dark grey and black evolutionary strata in the *Microbotryum violaceum melanantha* clade. Arrows correspond to the suppression of recombination steps. See supplementary Figure S5 for details on the steps. Box: Key to symbols in the figure.

The genealogies displayed by the genes ancestrally located in the three regions analysed, i.e. PR-to-centromere, PR short arm and HD-to-centromere, were incongruent for some nodes (Figure 4), in particular for the placement of alternate mating types in the *M. v. gracilicaulis - M. v. viscidula* - *M. v. melanantha* clade. We indeed detected trans-specific polymorphism for these three species (i.e., clustering of allele according to mating type rather than according to species) in the gene genealogies of the HD-to-centromere but not the PR-to-centromere ancestral regions. The genes ancestrally located in the PR-chromosome short arm displayed trans-specific polymorphism between *M. v. viscidula* and *M. v. melanantha*, but not shared with *M. v. gracilicaulis*, while there was no trans-specific polymorphism at all in this clade in the PR-to-centromere ancestral region. This suggests that full HD-PR linkage in this clade was achieved in several steps, some of which occurred independently in the different species (Figure 4 and Supplementary Figure S5). The chromosomal rearrangement and the suppression of recombination in the HD-to-centromere region have most likely occurred at the base of this three-species clade. Indeed, the ancestral recombination suppression in the HD-to-centromere region would not be beneficial unless the chromosomal rearrangement had already occurred, linking the HD and PR loci one to each other. The recombination suppression has then extended after the divergence of *M. v. gracilicaulis*, and then again further after the speciation between *M. v. viscidula* and *M. v. melanantha* (Figure 4A, 4B and 4C). This stepwise completion of recombination suppression led to the formation of three distinct evolutionary strata in the *M. v. viscidula-M. v. melanantha* clade, that we called light grey, dark grey and black strata (Figure 4E), and which can be also seen in the d_S_ plots (Figure 3B). The estimated dates of recombination suppression (Supplementary Table S5) indicated that the light grey stratum (HD-to-centromere) occurred in the *M. v. viscidula-M. v. melanantha* clade 1.5 MYA (in the same event as in the *M. v. gracilicaulis* lineage), while the dark grey stratum (PR-chromosome short arm) occurred 0.88 MYA and the black strata (PR-to-centromere) between 0.57 MYA in *M. v. melanantha* and 0.6 MYA in *M. v. viscidula*. In the *M. v. gracilicaulis* lineage, the black stratum (PR-chromosome short arm plus PR-to-centromere) evolved 0.8 to 0.9 MYA (Supplementary Figure S5).

In *M. v. lateriflora* parasitizing *Moehringia lateriflora* (Figure 2F) and *M. v. parryi* parazitizing *Silene parryi* (Supplementary Figure S4D), the mating-type chromosomes were formed by a fusion between the long arm of the PR chromosome and the short arm of the HD chromosome, following the same scenario as *M. v. caroliniana* and *M. tatarinowii* (Figure 1). The placement of *M. v. parryi* as a sister species of *M. v. caroliniana* in the phylogeny suggests that the mating-type locus linkage event could have occurred before the divergence of these two species, which was confirmed by the trans-specific polymorphism found between these two species in their black stratum (Figure 4). In contrast, the strongly supported placement of *M. v. lateriflora* within the clade formed by *M. lagerheimii* and *M. saponariae* (Figure 1), both with mating-type loci linked to their respective centromere, indicates that the mating-type chromosome rearrangements in *M. v. lateriflora* occurred independently from that in *M. v. caroliniana, M. v. parryi* and *M. v. tatarinowii*, representing convergence. The exact same breakpoints were involved in the convergent rearrangements, exactly at the centromere when we consider that the ancestral state of the PR chromosome corresponds to the gene order of *M. lagerheimii*.

We found a large range for the estimated dates of recombination suppression events between mating-type loci, from 0.15 MYA in *M. silenes-acaulis* to 3.58 MYA *M. scabiosae* (Figure 4, Supplementary Table S5). In *M. scabiosae*, the much older dates of recombination suppression in the HD-to-centromere region than in the PR-to-centromere region suggest that, in this lineage too, the recombination suppression between the two mating-type loci has likely occurred in two distinct evolutionary strata, which can also be observed in the d_S_ plot previously published (Branco et al 20218). Actually, the estimated dates of recombination suppression were more ancient for the HD-to-centromere region than in the long arm of the PR chromosome in most species (Supplementary Table S5); this suggests that recombination was not completely suppressed all along the region between the two mating-type loci immediately after the chromosomal rearrangements in the various species, but extended progressively from the mating-type loci until joining to complete recombination cessation.

### Evolutionary strata beyond mating-type loci

In order to investigate the existence of young evolutionary strata, we inspected the d_S_ values between the alleles in a_1_ and a_2_ genomes along the mating-type chromosomes according to the ancestral gene order inferred from *M. lagerheimii*. In all the four species with new genome sequences, d_S_ was high between alleles at the genes located ancestrally between the mating-type loci, in agreement with the complete recombination suppression linking mating-type loci (Figures 3B, C, D and E). As in previous studies (Branco et al., 2018, 2017), we called the regions without recombination linking the HD and PR loci in each species the black strata; note however that they do not all have the same origin or gene content (Figure 1). In *M. v. lateriflora*, the moderate level of rearrangements (Figure 2C), together with the low level of differentiation between the genes of the black strata (Figure 3D), confirm that the mating-type locus linkage event occurred very recently (date estimate 0.19 to 0.41 MYA, Supplementary Table S5). More extensive rearrangements between a_1_ and a_2_ mating type chromosomes in *M. v. gracilicaulis, M. v. viscidula* and *M. v. parryi* (Figure 2C; Supplementary Figures S1A and S1C), as well as higher d_S_ values in their black strata (Figures 3B, C and E), confirm the older linkage of the mating-type loci in these species (Supplementary Table S5).

We found, in all the species studied here, the footprints of the same ancient shared evolutionary strata as in other *Microbotryum* species, with high d_S_ levels in the blue and purple regions in proximity to the HD and PR loci, respectively (Figure 3), thus likely representing stepwise extensions of recombination suppression beyond mating-type loci at the basis of the *Microbotryum* clade (Figure 1). These different evolutionary strata were not initially delimited only based on their d_S_ levels, but by i) the set of species displaying non-zero d_S_ values in these genomic regions and ii) the level of trans-specific polymorphism, two strong indicators of the origin of the strata in the phylogeny. Further extension of the recombination suppression beyond the PR locus and its ancient purple stratum was detected in *M. v. gracilicaulis* and *M. v. viscidula* (Figures 3B and 3C), corresponding to the previously identified orange stratum in the *M. lychnidis-dioicae* clade as well as in *M. silenes-acaulis* and *M. v. paradoxa* (Figure 1). In agreement with the previous inference that the orange stratum evolved after the divergence of the *M. lagerheimii* - *M. saponariae* clade (Hartmann et al., 2021), we did not find traces of the orange stratum in *M. v. lateriflora* or *M. v. parryi* (Figure 3C and 3D). However, the orange stratum seemed to be present in *M. v. tatarinowii* (Fig. 1; Carpentier et al. 2021), which suggests an independent evolution in this later lineage. We did not find on *M. v. parryi* footprints of the light blue stratum previously identified in *M. v. caroliniana* (Branco et al. 2018), neither from the d_S_ plot or in the trans-specific polymorphism analyses, supporting the inference that this stratum evolved very recently in *M. v. caroliniana*, later than the black stratum that is shared with *M. v. parryi* (trans-specific black stratum date 2.23-2.79 MYA, *M. v. caroliniana* light blue stratum date 0.26 MYA, speciation date 1.42 MYA, Figure 4 and Supplementary Table S5).

## Discussion

In the present study, we obtained high-quality assemblies of alternative mating types for four additional *Microbotryum* anther-smut fungi, in addition to the 13 already available. We found additional events of independent chromosomal rearrangements bringing the two mating-type loci onto the same chromosome followed by recombination suppression linking the two mating-type loci to each other. In total, this makes at least nine independent and convergent events of mating-type locus linkage across the *Microbotryum* genus. The dates of recombination suppression between the two mating-type loci ranged from 0.15 to 3.58 million years ago. In addition, we found further strong support for independent events of recombination suppression between mating-type loci and centromeres in two *Microbotryum* species, through the phylogenetic placement of an additional species whose genomes we sequenced here. With these four events of recombination suppression (two species times two centromere-mating-type locus linkage events), this makes a total of 13 events of recombination suppression involving mating-type loci across the phylogeny of *Microbotryum* anther-smut fungi, and even more if we count the independent completion events of recombination cessation in the *M. v. melanantha* clade. As the *Microbotryum* genus likely contains more than a hundred species (Hood et al., 2010; Lutz et al., 2008), these results suggest that there remains a rich resource of genomic diversity in the evolution of suppressed recombination in linkage to reproductive compatibility loci. Given the high number of convergent events having linked the two mating-type loci on the same chromosome, one may wonder why it did not occur earlier, at the basis of the *Microbotryum* clade. A speculative answer is that similar chromosomal rearrangements need to occur on the two mating-type chromosomes to avoid unbalanced meiosis (Fraser et al., 2004), which may not occur that easily, even if the number of convergent events across the phylogeny show that it can happen repeatedly.

Additional independent events of mating-type locus linkage also occurred in other basidiomycete genera, e.g., *Ustilago* and *Cryptococcus* (Bakkeren and Kronstad, 1994; Hartmann et al., 2021; Sun et al., 2019). Such striking and repeated convergence shows that strong selection can lead to similar evolution repeatedly. The linkage of HD and PR loci has likely been selected for increasing gamete compatibility odds under selfing (Branco et al 2017, Hood 2002, 2004). With a single locus and two alleles, gametes are indeed compatible with 50% of other gametes, while the odd is only 25% with two loci and two alleles each, as fungi can only mate when alleles are different at both loci. Linkage between each of the two mating-type loci and their respective centromere gives the same odds of gamete compatibility as mating-type loci linkage, but only within meiotic tetrads, not among tetrads of a given diploid individual where the odds fall to 25%. This may be why only two *Microbotryum* species evolved PR- and HD-centromere linkage without chromosome fusion (Carpentier et al. 2019) whereas HD-PR mating-type loci linkage evolved a dozen times. Such two-fold increase in the odds of compatibility may be important to maximize the chances of plant infection, as a higher number of dikaryons on a given plant increases disease probability (Kaltz et al., 1999; Roche et al., 1995), and it may give a time advantage under competition situations, as a single genotype typically has a resident advantage at parasitizing a plant even when several genotypes are deposited on the plant (Day, 1980; Gibson et al., 2012; Giraud et al., 2005; Hood, 2003; López-Villavicencio et al., 2007). The repeated convergent evolution of mating-type locus linkage shows that natural selection is very powerful in the face of contingency (i.e. the stochasticity associated with initial conditions and mutations occurring randomly) and can make evolution repeatable.

We did not find new types of chromosomal rearrangements having led to HD and PR on a single chromosome, and instead found repeated independent evolution of the same four types of chromosome fusion/fission events, with breakpoints at centromeres. This suggests that the convergence events of HD-PR linkage have occurred by convergence of a limited number of chromosomal rearrangement types and that rearrangements at centromeres are more likely than at other places in chromosomes. Such rearrangements at centromeres have also been reported in the other basidiomycetes having undergone HD-PR locus linkage (Sun et al., 2017).

In the *M. v. gracilicaulis-M. v. viscidula-M. v. melanantha* clade, we found that recombination suppression in the HD-to-centromere region was ancestral to the clade while the recombination suppression in the PR short arm was shared only by *M. v. viscidula* and *M. v. melanantha*, and the the recombination suppression in the PR-to-centromere region occurred independently in each species, completing the PR-HD linkage. Recombination suppression between the HD locus and its centromere without linkage to PR would bring no advantage in terms of odds of gamete compatibility; the chromosomal rearrangement that has brought the HD and PR loci on the same chromosome is therefore likely ancestral to the clade, which is supported by the trans-specific polymorphism found at some genes in the short arm of the PR chromosome. There would have been initially only partial linkage between HD and PR if recombination was allowed in the PR-to-centromere region, but the increased rates of gamete compatibility would still have been beneficial compared to full independence of mating-type loci. Later, completion of recombination cessation would also have been a beneficial step, further increasing rates of gamete compatibility. We also found evidence for gradual completion of recombination cessation in the *M. v. caroliniana* - *M. v. parryi* clade.

In conclusion, our findings further support the anther-smut fungi as excellent models to study the evolution of recombination suppression and its consequences, providing many independent cases of recombination cessation across a wide range of ages.

## Data access

Sequencing data and genome assemblies were deposited at GenBank under the BioProject PRJNA771266.

## Acknowledgements

This work was supported by the National Institute of Health (NIH) grant R15GM119092 to M. E. H., the Louis D. Foundation award and EvolSexChrom ERC advanced grant #832352 to T. G., and an IDEX Paris-Saclay Bochum University PhD grant to M.D. We thank Kurt Hasselman, John Bain and Hui Tang who collected the strains (with an avoided encounter with a grizzly bear for John Bain). We thank Cécile Fairhead for help with DNA extraction. PacBio sequencing was conducted at the IGM Genomics Center, University of California, San Diego, La Jolla, California

## Author contributions

T.G., R.C.R.d.l.V. and M.E.H designed and supervised the study. T.G., M.E.H, D.B. and M.D. contributed to get funding. F.C., R.C.R.d.l.V. and M.E.H. obtained the genome assemblies. M.D., F.C., R.C.R.d.l.V. performed the genomic analyses. T.G., R.C.R.d.l.V. and M.D. wrote the original draft. All authors contributed to the manuscript.

## Figures

**Supplementary figure S1:**
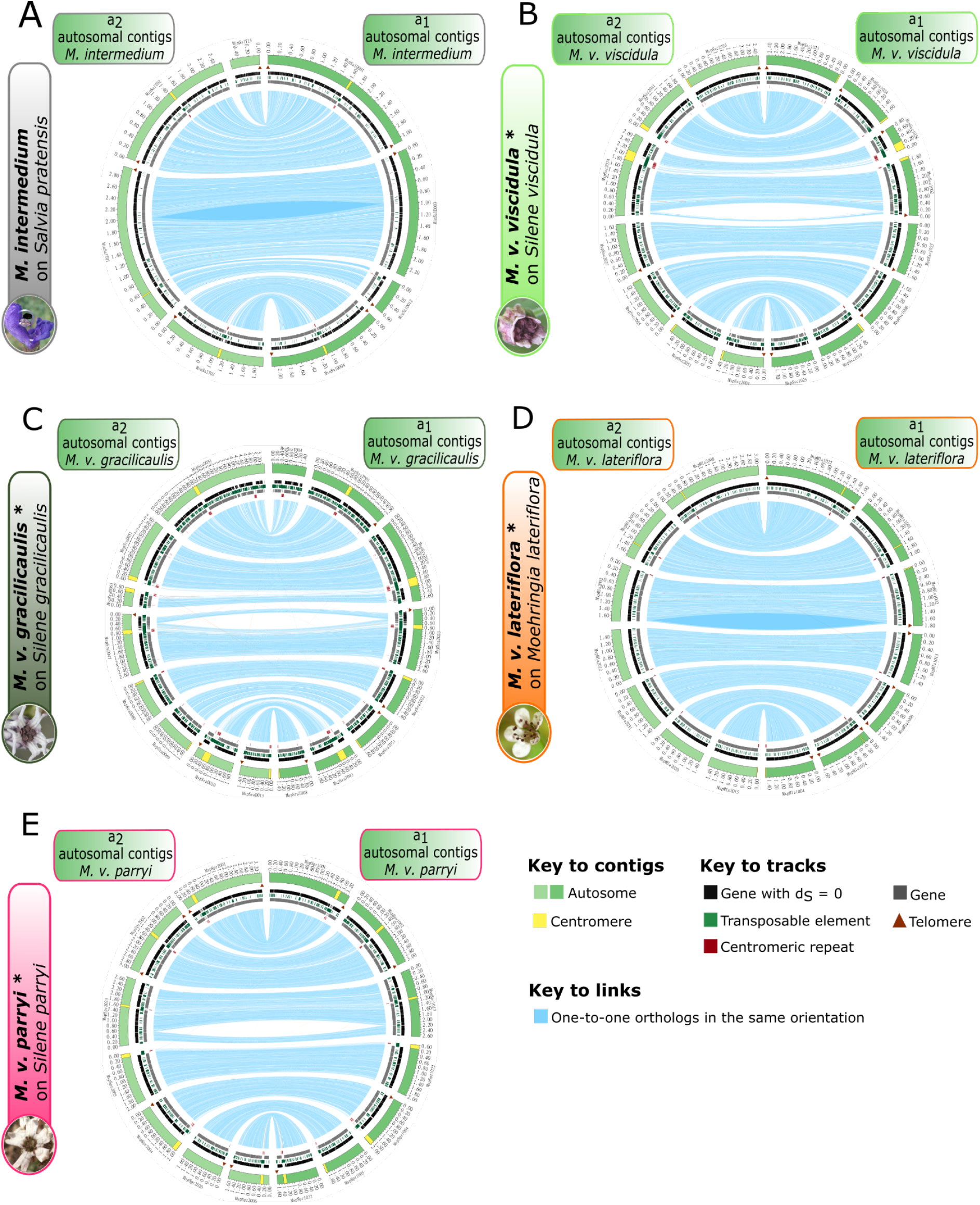
Comparison of gene order between haploid genomes of opposite mating types for the largest autosomal contigs in *Microbotryum intermedium, M. violaceum viscidula, M. v. gracilicaulis, M. v. lateriflora* and *M. v. parryi*. Comparison of gene orders between a_1_ and a_2_ haploid genomes for autosomal contigs in A) *M. intermedium*, B) *M. v. viscidula*, C) *M. v. gracilicaulis*, D) *M. v. lateriflora* and D) *M. v. parryi*. The outer track represents contigs, with length ticks every 200 kb. Contigs of the haploid genomes of a_1_ and a_2_ mating types are depicted in light green and green, respectively. The centromeres are represented in yellow. The telomeres are indicated by brown triangles. Black marks on the inner track represent the genes with null synonymous divergence between a_1_ and a_2_ alleles. Green marks on the inner track represent transposable element (TE) copies. Gray marks on the inner track correspond to genes. Red marks on the inner track correspond to the putative centromeric repeats. The link width is proportional to the corresponding gene length. Blue links link alleles in the same orientation.

**Supplementary figure S2:**
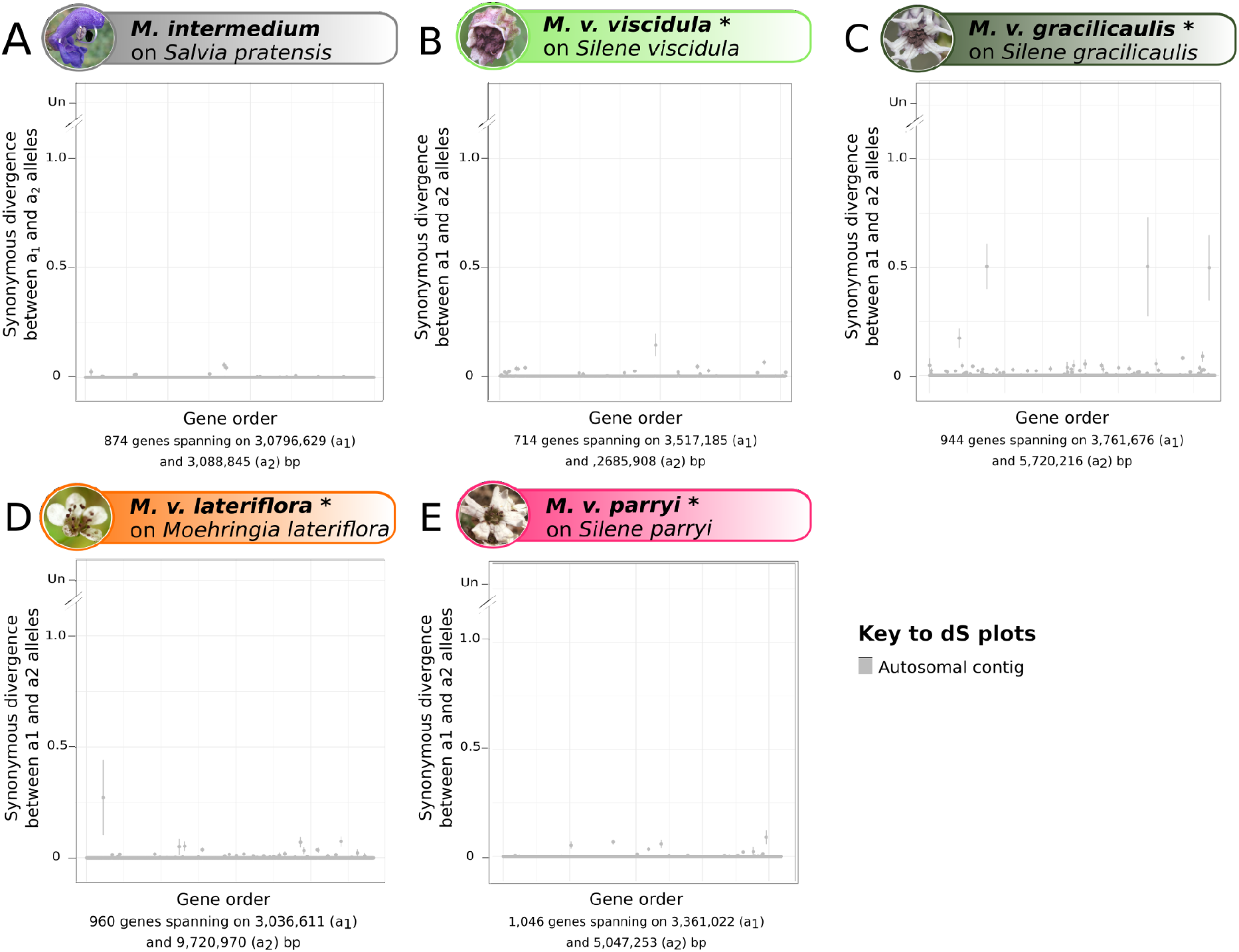
Differentiation between alleles of haploid genomes isolated from a diploid individual, along autosomal contigs in *Microbotryum* fungi. Per-gene synonymous divergence between alleles present in the haploid a_1_ and a_2_ genomes from a given diploid individual, along the largest autosomal contigs of each of A) *M. intermedium*, B) *M. v. viscidula*, C) *M. v. gracilicaulis*, D) *M. v. lateriflora* and E) *M. v. parryi*, plotted along the chromosome. The number of genes and average of the a_1_ and a_2_ contig lengths are indicated on the X axis.

**Supplementary figure S3:**
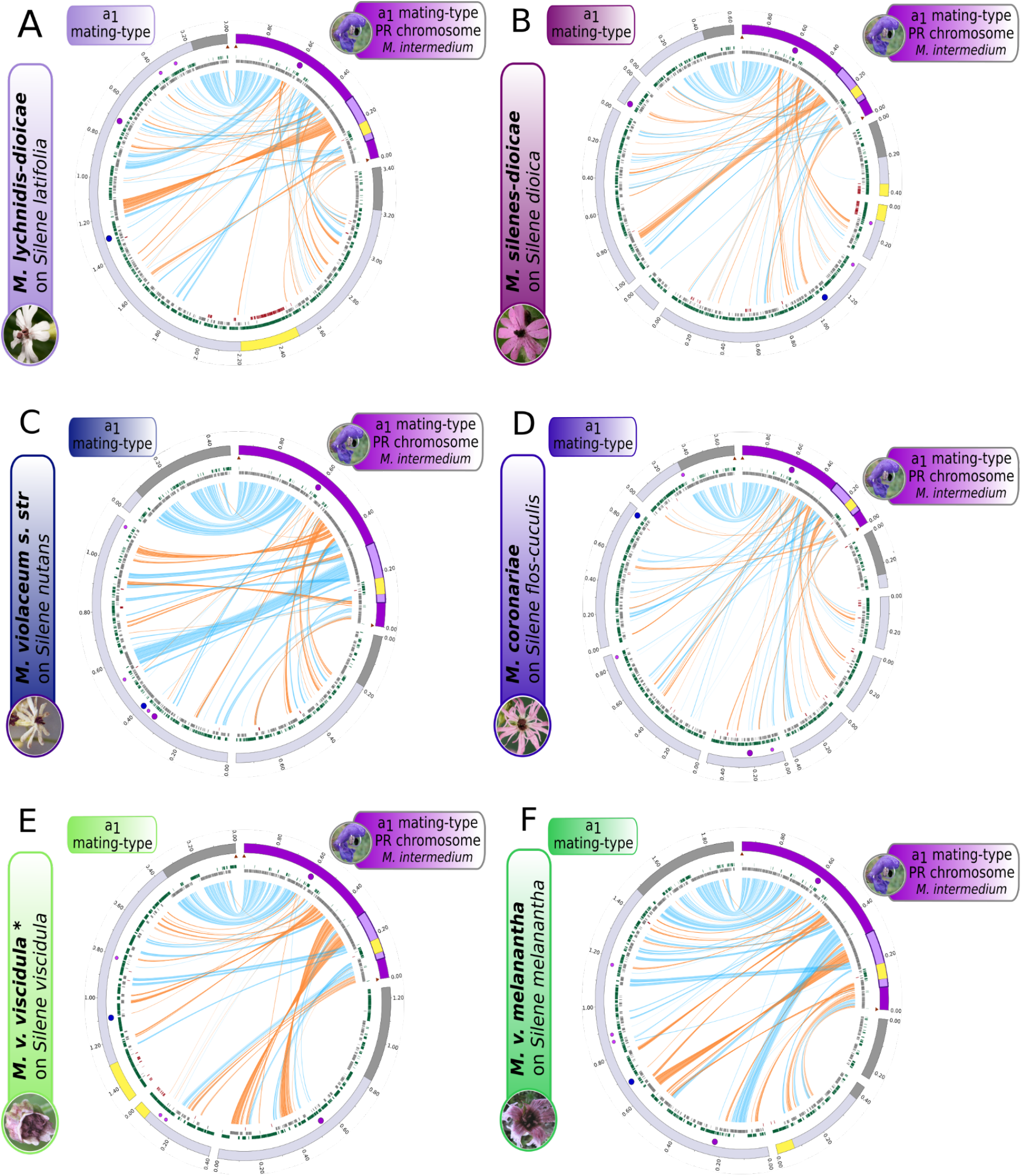
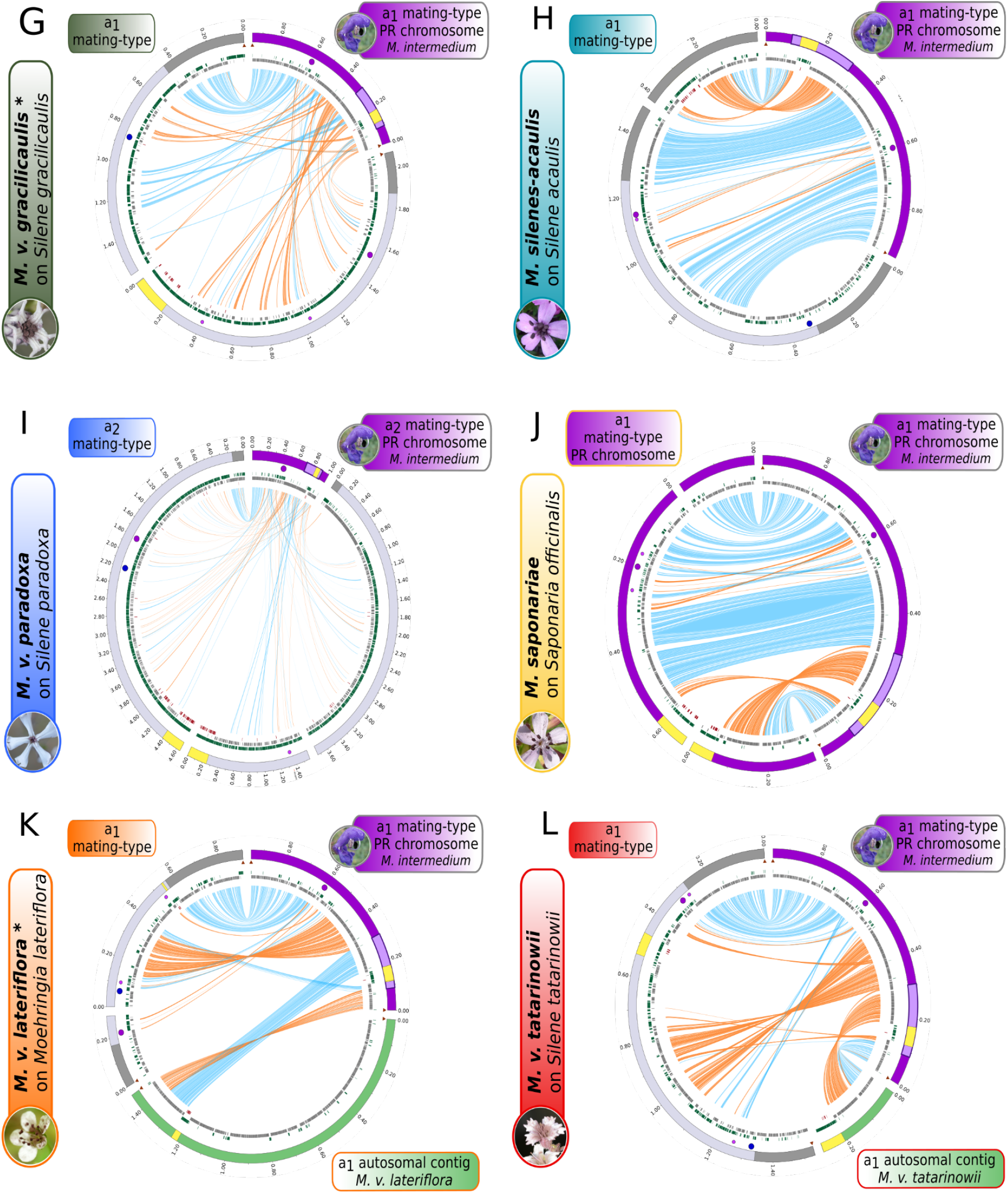
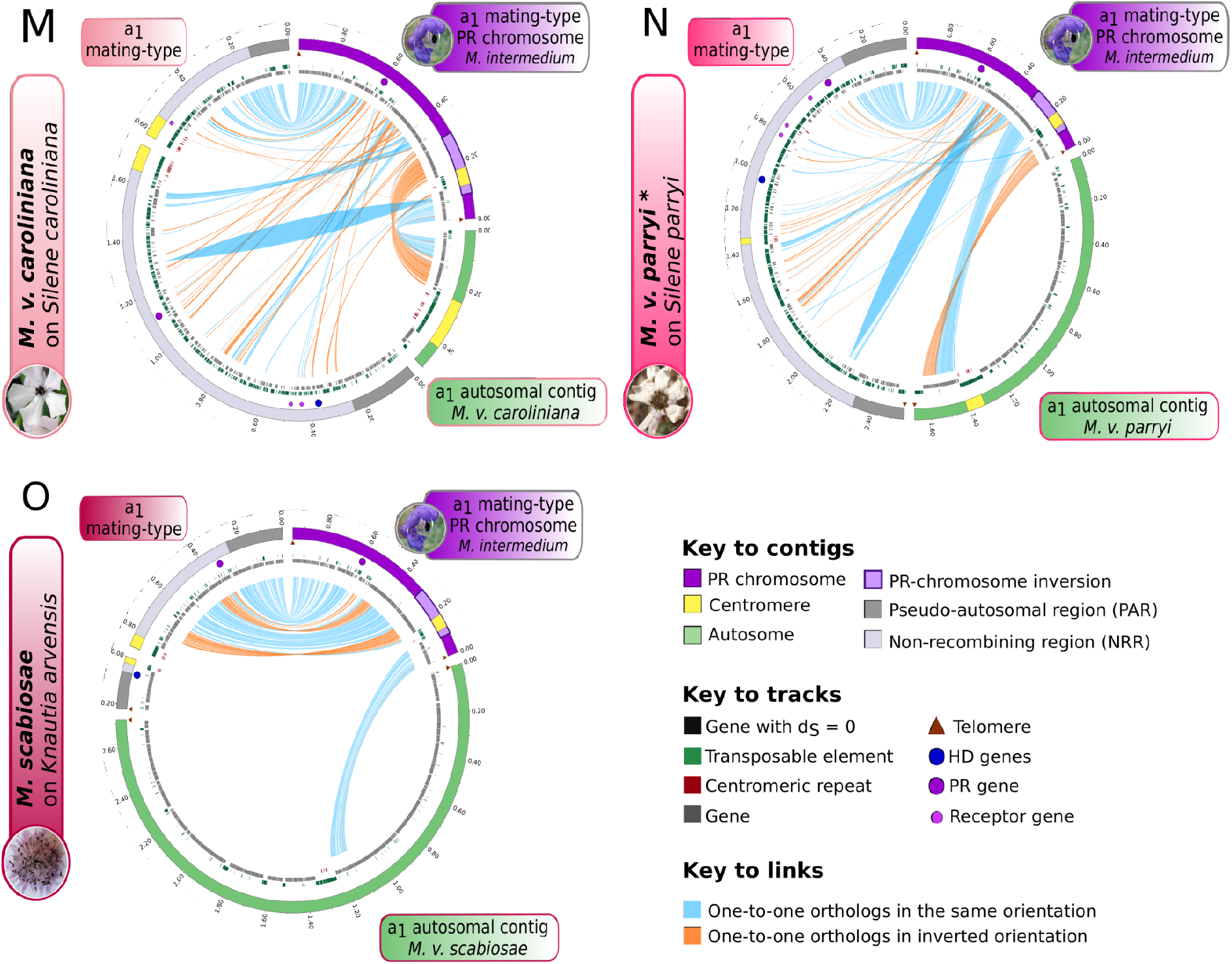
Inter-specific comparison of gene order between the PR mating-type chromosome of *Microbotryum intermedium* and the mating-type chromosomes of all the species with suppressed recombination. Comparison of gene order between a_1_ mating-type PR chromosome of *M. intermedium* and their homologues on the PR-chromosome of A) *M. lychnidis-dioicae*, B) *M. silenes-dioicae*, C) *M. violaceum s. str*., D) *M. coronariae*, E) *M. v. viscidula*, F) *M. v. melanantha*, G) *M. v. gracilicaulis*, H) *M. silenes-acaulis*, I) *M. v. paradoxa*, J) *M. saponariae*, K) *M. v. lateriflora*, L) *M. v. tatarinowii*, M) *M. v. caroliniana*, N) *M. v. parryi* and O) *M. scabiosae*. The PR, HD and pheromone genes are represented by purple, blue and small light purple circles, respectively. The outer track represents contigs, with external ticks every 200 kb. The PR mating-type chromosome of *M. intermedium* is represented in purple. (A to O). The region of the inversion having occurred after divergence with *M. scabiosae* (Fig. 1), in the region encompassing the centromere of the PR chromosome, is highlighted on the outer track in light purple. The non-recombining region (NRR) and the pseudo-autosomal region (PAR) of the mating-type chromosomes of *M. lychnidis-dioicae* (A), *M. silenes-dioicae* (B), *M. coronariae* (C), *M. violaceum s. str*. (D), *M. v. viscidula* (E), *M. v. melanantha* (F), *M. v. gracilicaulis* (G), *M. silenes-acaulis* (H), *M. v. paradoxa* (I), *M. saponariae* (J), *M. v. lateriflora* (K), *M. v. tatarinowii* (L), *M. v. caroliniana* (M), *M. v. parryi* (N) and *M. scabiosae* (O)are represented in light and dark gray, respectively. Green contigs correspond to autosomal contigs, in *M. v. lateriflora* (K), *M. v. tatarinowii* (L), *M. v. caroliniana* (M), *M. v. parryi* (N) and *M. scabiosae* (O). The centromeres are represented in yellow. The telomeres are indicated by brown triangles. Black marks on the inner track represent the genes with null synonymous divergence between a_1_ and a_2_ alleles. Green marks on the inner track represent transposable element (TE) copies. Gray marks on the inner track correspond to genes. Red marks on the inner track correspond to the putative centromeric repeats. Blue and orange lines link alleles, the latter corresponding to inversions. The link width is proportional to the corresponding gene length.

**Supplementary figure S4:**
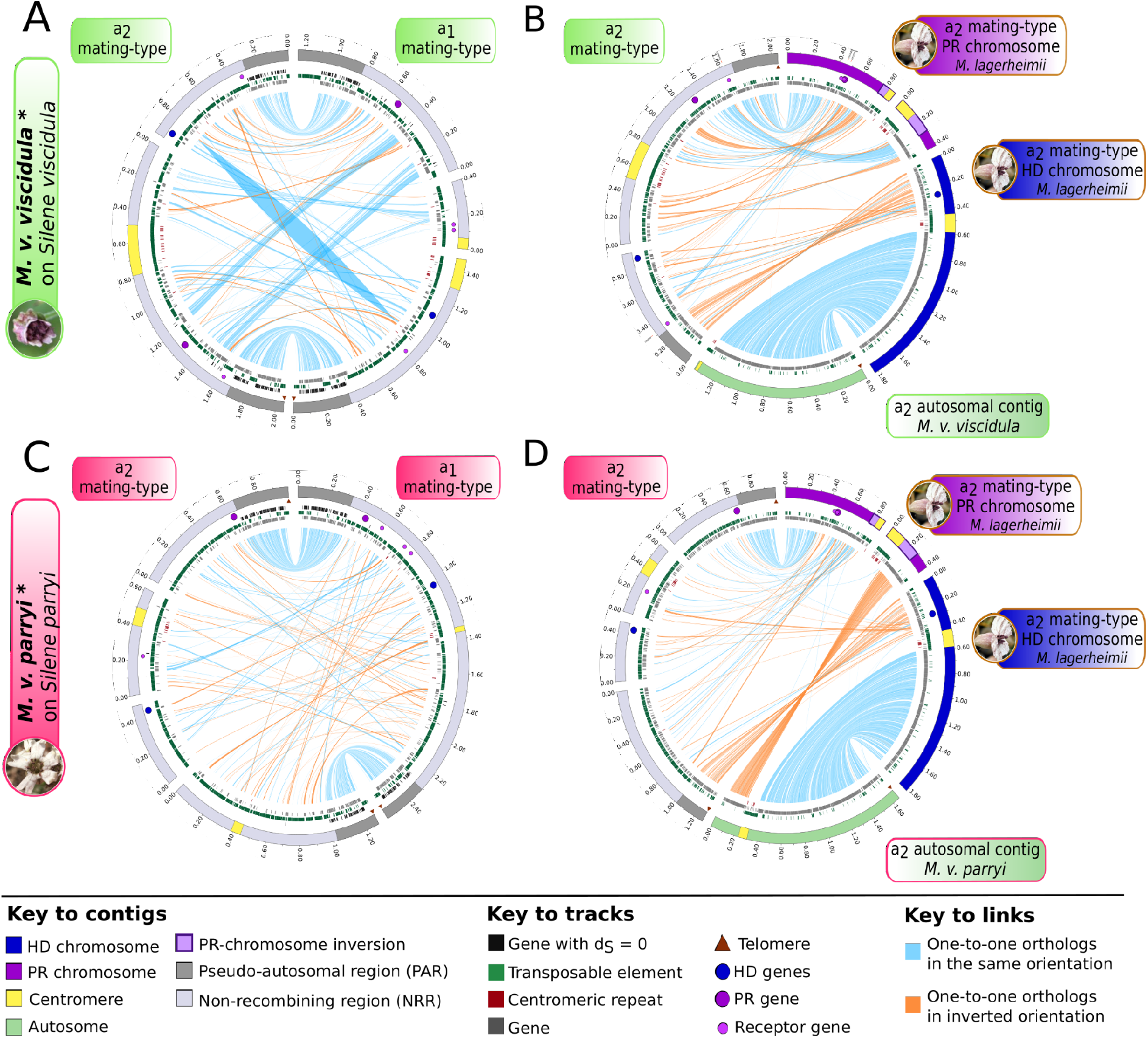
Inter- and intra-specific comparison of gene order between mating-type chromosomes in *Microbotryum violaceum viscidula* and *M. v. parryi*. Synteny plots between A) a_1_ and a_2_ mating-type chromosomes of *M. v. viscidula*, B) a_2_ mating-type chromosome of *M. lagerheimii* and their homologues in *M. v. viscidula*, C) a_1_ and a_2_ mating-type chromosomes of *M. v. parryi*, D) a_2_ mating-type chromosome of *M. intermedium* and their homologues in *M. v. parryi*. Comparisons of *M. lagerheimii* a_2_ HD and PR chromosomes and *M. v. viscidula* (B) and *M. v. parryi* (F) a_2_ mating-type contigs to infer chromosomal rearrangements having led to HD-PR linkage, considering *M. lagerheimii* chromosomes as a proxy for the ancestral chromosomal state. The PR, HD and pheromone genes are represented by purple, blue and small light purple circles, respectively. The outer track represents contigs, with length ticks every 200 kb. The HD and PR mating-type chromosomes of *M. lagerheimii* are represented in blue and purple, respectively (B and D). The region of inversion having occurred after divergence with *M. scabiosae* (Fig. 1), in the region encompassing the centromere of the PR chromosome, is highlighted in light purple. The non-recombining region (NRR) and the pseudo-autosomal region (PAR) of the mating-type chromosomes of *M. v. viscidula* (A and B) and *M. v. parryi* (C and D) are represented in light and dark gray, respectively. Green contigs correspond to autosomal contigs, in *M. v. viscidula* (B) and *M. v. parryi* (D). The centromeres are represented in yellow. The telomeres are indicated by brown triangles. Black marks on the inner track represent the genes with null synonymous divergence between a_1_ and a_2_ alleles. Green marks on the inner track represent transposable element (TE) copies. Gray marks on the inner track correspond to genes. Red marks on the inner track correspond to the putative centromeric repeats. Blue and orange lines link alleles, the latter corresponding to inversions. The link width is proportional to the corresponding gene length.

**Supplementary figure S5:**
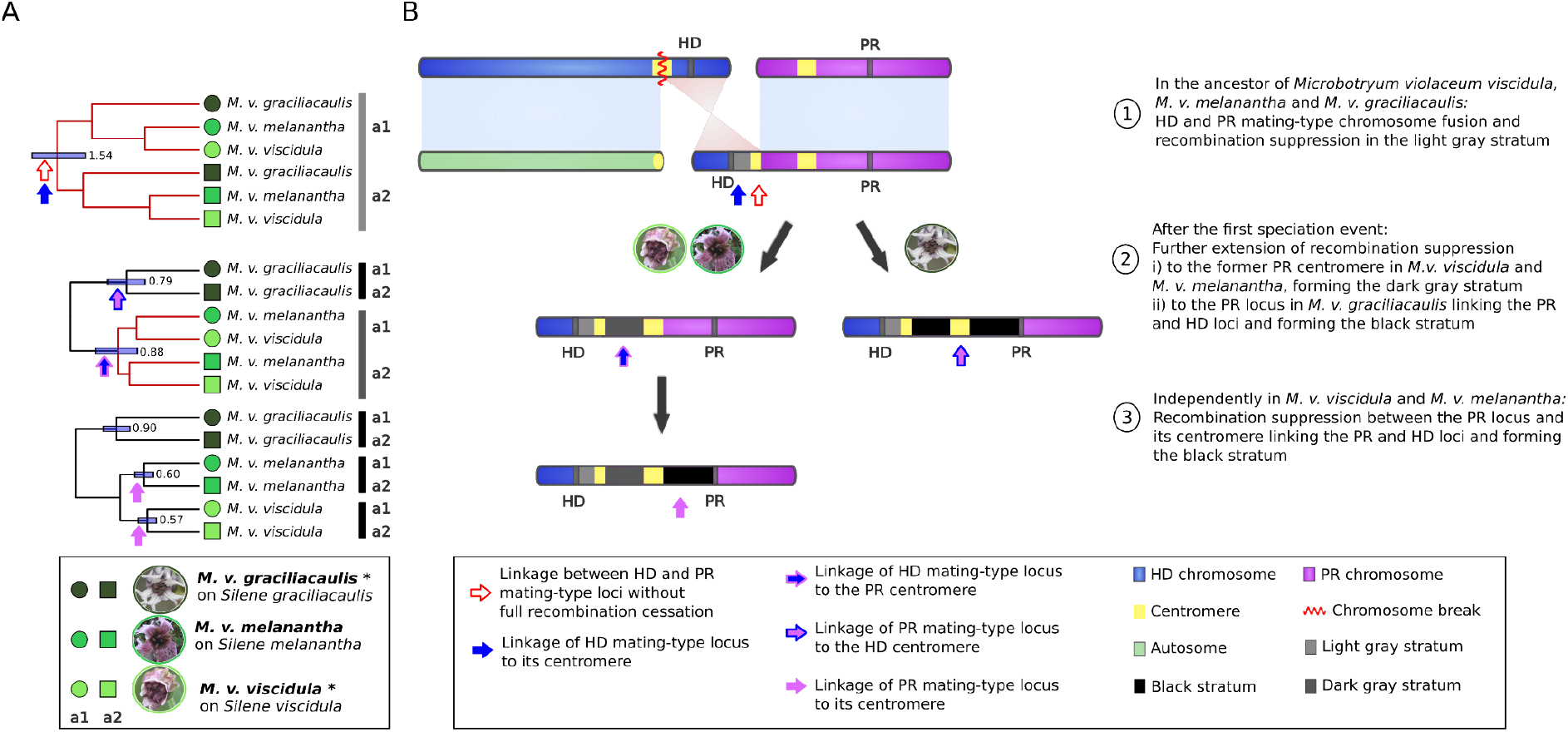
Reconstructed scenario of stepwise recombination suppression between the HD and PR mating-type loci in the *Microbotryum violaceum melanantha* clade. A) Timetree of conserved genes in the regions between the HD locus and its ancestral centromere (upper panel), between the short arm telomere and the centromere in the ancestral PR chromosomes (middle panel) or between the PR locus and its ancestral centromere (bottom panel). Topologies are significantly different, showing full trans-specific polymorphism between a_1_ and a_2_ alleles (light gray stratum, upper panel), trans-specific polymorphism between a_1_ and a_2_ alleles of *M. v. melanantha* and *M. v. viscidula* (dark grey stratum, middle panel) or no trans-specific polymorphism (bottom panel). Trans-specific polymorphisms are indicated by red branches. B) Reconstructed stepwise recombination suppression having generated the light gray, dark grey and black evolutionary strata. Arrows correspond to the suppression of recombination steps

